# Integrated small and long RNA sequencing in single oocytes reveals piRNA-mediated transposon repression during human oogenesis

**DOI:** 10.1101/2025.07.22.666229

**Authors:** Fengjuan Zhang, Hongdao Zhang, Yali xiao, Miao Liu, Aimin Ren, Suying Liu, Ligang Wu

## Abstract

The piwi-interacting RNA (piRNA) pathway plays a pivotal role in controlling transposable element (TE) activity, which is crucial for the developmental competence of gametogenesis. Although piRNAs have been studied in golden hamsters and other representative mammals, little is known about the relationship between distinct piRNA populations and their regulatory effects on TEs in human oocytes. In this study, we simultaneously profiled small and long RNA transcriptomes in individual human oocytes across four developmental stages. piRNAs, especially PIWIL3-associated short piRNAs (short-piRNAs), are the predominant small non-coding RNAs during human oogenesis. A marked increase in short-piRNAs after the primordial follicle stage coincided with a global downregulation of TE expression, particularly LINE-1 (L1) and endogenous retroviruses (ERVs). On the other hand, PIWIL1- and PIWIL2-associated long piRNAs (long-piRNAs) were correlated with the silencing of certain specific ERV subfamilies. Genomic-context analyses revealed that highly productive piRNA clusters have evolved asymmetric antisense insertion bias toward L1 and ERVs, contributing to TE families-specific regulation. Our findings highlight the global effect of piRNA-mediated TEs repression, with short-piRNAs acting as the primary and broad-spectrum suppressors, and long-piRNAs providing coordinated ERV-specific silencing. In summary, this study provides a valuable dataset of small and long RNA co-expression landscapes in developing human oocytes and offers insights into the coordinated yet distinct roles of different PIWI/piRNA classes in repressing TEs during human oogenesis.

## Introduction

Transposable elements (TEs) are mobile genetic elements that are capable of invading and propagating within host genomes. While TE activity has played a major role in shaping genome evolution, their mobilization also poses a threat to genomic integrity by inducing DNA double-strand breaks and facilitating illegitimate recombination^1, 2^. Based on their mobilization mechanisms, TEs are categorized into two major classes^3,4^. Class I elements, or retrotransposons, mobilize via an RNA intermediate and include both long terminal repeat (LTR) and non-LTR retrotransposons. LTR retrotransposons primarily consist of endogenous retroviruses (ERVs) and ERV-derived solo LTR elements. Non-LTR retrotransposons comprise long interspersed nuclear elements (LINEs) such as *L1*, *CR1*, and *L2*; short interspersed nuclear elements (SINEs) including *SINE1/7SL*, *SINE2/tRNA*, and *SINE3/5S rRNA*; and a composite element SINE-VNTR-Alu (*SVA*). Class II elements, or DNA transposons, mobilize via a DNA intermediate. Although the vast majority of TEs, all DNA transposons included, are extinct, several retrotransposon families remain active in the human population^5^. These active elements can mobilize autonomously or rely on trans-acting factors encoded by other autonomous elements. Notably, subfamilies such as *L1PA1, SVA,* and *AluY* exhibit insertional polymorphisms among human individuals, reflecting ongoing transpositional activity^6, 7^. Among ERVs*, HERV-*K is the only lineage known to exhibit polymorphic insertions in modern humans^8, 9^. Additional support for their ongoing activity comes from somatic insertions observed in neuronal and cancer cells^10-15^, indicating their potential to mobilize post-zygotically and contribute to somatic mosaicism.

While TE repression is effectively maintained in somatic cells, genome-wide epigenetic reprogramming during primordial germ cell formation erases most DNA methylation marks, creating a vulnerable window for TE activation^16-18^. In females, this epigenetic state persists until the initiation of follicle growth^19^. Unchecked TE activity during this critical period can lead to genomic instability, meiotic arrest, and infertility^20,21^. The piRNA pathway is a major defense mechanism against TE activity in the germline. PIWI proteins, a subfamily of ARGONAUTE, associate with piRNAs to mediate TE suppression via both post-transcriptional cleavage and epigenetic silencing^22-31^. In *Drosophila* and *zebrafish*, disruption of PIWI proteins leads to sterility in both males and females due to uncontrolled TE activity^22, 32-36^. In mice, the loss of *PIWIL1 (MIWI)*, *PIWIL2 (MILI)*, or *PIWIL4 (MIWI2)* results in TE dysregulation and male infertility, whereas female oogenesis remains largely unaffected^24, 36-38^. Recent transcriptomic analyses of mammalian oocytes have revealed substantial variability in small RNA composition across species^39^. Interestingly, mouse oocytes exclusively express endo-siRNAs rather than piRNAs, which are processed by a unique Dicer isoform (DicerO) and functionally substitute for piRNAs in TE suppression^40^. In contrast, human and most other mammalian oocytes predominantly express PIWIL3-associated short piRNAs (short-piRNAs), which account for most of the small RNAs in these cells^39, 41, 42^. Functional studies using golden hamsters as a model have demonstrated that loss of the piRNA pathway compromises oocyte competence and leads to early embryonic arrest^43-45^. Additionally, knockout studies of individual PIWI proteins suggest their non-redundant roles in TE suppression and gametogenesis^46^.

However, the temporal dynamics of piRNAs expression during human oogenesis and the functional relationship between distinct piRNA populations and TE regulation remain poorly understood. To address this gap, we profiled both small and long RNA transcriptomes in individual human oocytes across four key developmental stages. Our analyses reveal a global repression of TEs following the primary follicle stage, with short-piRNAs serving as the predominant and broad suppressors and long-piRNAs providing coordinated, ERV-specific repression. These findings highlight the distinct yet coordinated roles of piRNA populations in TE silencing and provide new insights into the regulatory landscape of human oocyte development.

## Results

### Profiling small and long transcripts in developing human oocytes

A total of 30 human oocytes spanning four developmental stages, primordial follicle (PrF), primary follicle (PF), germinal vesicle (GV), and metaphase II (MII), were collected. Small and long RNA transcriptomes were simultaneously profiled at the single-oocyte level using our previously established protocol^39, 43, 46^ (Fig. 1a, *Materials* and *Methods*). Following quality control filtering, 23 oocytes were retained for further analyses, comprising 5 PrF, 9 PF, 4 GV, and 5 MII stage oocytes (Supplementary Data 1). On average, 5.88 million raw reads per small RNA library and 7.66 million reads per long RNA library were obtained (Supplementary Data 2). Among these retained oocytes, an average of 87.88% of small RNA-seq reads were deemed usable, with a genome mapping rate of 71.17%. Long RNA-seq libraries showed mapping rates ranging from 87.84% to 95.38%. Across all oocytes, we detected an average of 117 miRNAs, 113,861 piRNAs, and 16,039 genes (Supplementary Data 2).

**Figure 1.**
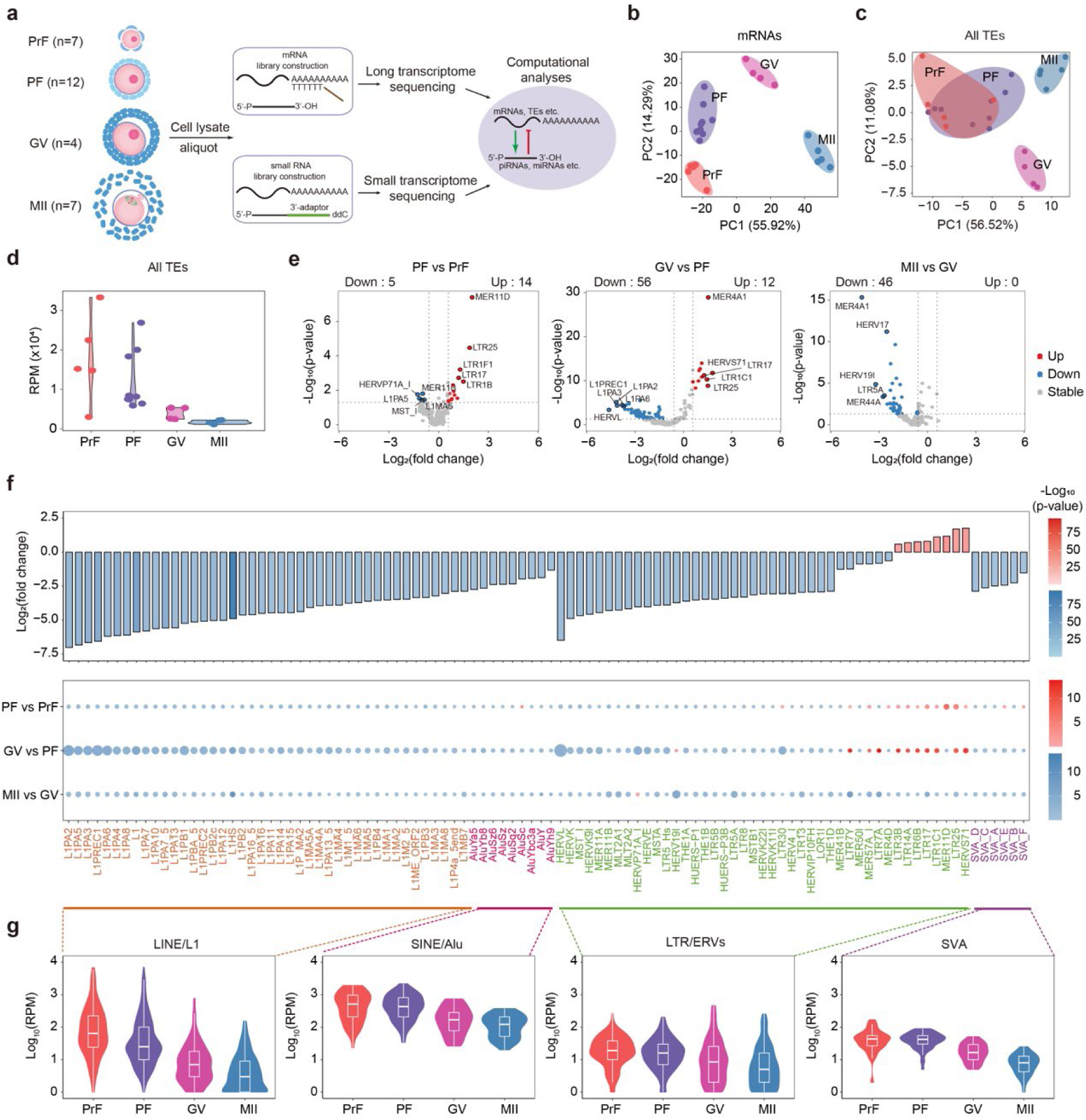
Global repression of TEs occurs after the primary follicle stages during human oocyte development and maturation. **a,** Schematic of the experimental strategy for simultaneous profiling of small and long RNA transcriptomes from single human oocytes at four developmental stages: primordial follicle (PrF), primary follicle (PF), germinal vesicle (GV), and metaphase II (MII). The number of oocytes collected at each stage is indicated. **b,** Principal component analysis (PCA) of protein-coding gene expression in individual oocytes across developmental stages. Expression levels were normalized to fragments per kilobase per million (FPKM). **c,** PCA of representative TE subfamily expression across oocyte development. TE expression was normalized to reads per million (RPM). **d**, Violin plots showing a progressive decline in the total TE expression (RPM) across developmental stages, indicating global TE repression during human oogenesis. **e,** Volcano plots of differentially expressed TE subfamilies across adjacent developmental stages (PF vs PrF, GV vs PF, MII vs GV). TE subfamilies with fold change ≥1.5 or ≤0.67 and p-value < 0.05 (Wald test) were considered significant. Numbers of upregulated (top right) and downregulated (top left) TE subfamilies are indicated. The top 5 most significant altered subfamilies in each comparison are labeled. **f,** *Upper Panel*: Bar plot showing log_2_-transformed fold changes and p-values of the TE subfamilies differentially expressed between the PrF and MII stages. *Lower Panel*: Fold changes and p-values of the same TE subfamilies across intermediate stages (PF vs Prf, GV vs PF, MII vs GV). Circle size reflects the magnitude of fold change; color intensity reflects -log_10_(p-value). Dashed lines group TE subfamilies by superfamily. **g,** Violin and box plots showing expression dynamics of differentially expressed TE subfamilies from the PrF to MII stages.

### Global downregulation pattern of TE during human oogenesis

Unsupervised principal component analysis (PCA) of protein-coding genes expression successfully segregated oocytes into four clusters, each corresponding to a different developmental stage (Fig. 1b and Supplementary Data 3). Gene Ontology (GO) enrichment analysis of stage-specific feature genes (558 for PrF, 570 for PF, 356 for GV, and 738 for MII) revealed key biological processes associated with each stage (Supplementary Fig. 1). Genes related to RNA and protein biosynthesis were significantly overrepresented in PrF and PF stages, while pathways involved in mitotic cell cycle regulation and nuclear division were enriched in GV and MII stages, reflecting ongoing oocyte growth and maturation^47^.

To assess TE expression dynamics during oogenesis, we mapped long RNA-seq reads to representative human TE sequences. PCA of TE expression revealed that PrF and PF oocytes clustered together, whereas GV and MII oocytes formed separate groups (Fig. 1c and Supplementary Data 4). A progressive decline in TE-derived transcript levels was observed across developmental stages, with a marked reduction occurring after the PF stage (Fig. 1d). Specifically, 56 TE subfamilies were significantly downregulated between the PF and GV stages, while 12 subfamilies were upregulated (Fig. 1e and Supplementary Data 5). From the GV to MII stages, most TE subfamilies exhibited reductions in expression, although this reduction was less pronounced compared to the PF-to-GV transition (Fig. 1e, f).

L1 experienced the most substantial repression during oocyte development, with an average 34.93-fold reduction from the PrF to MII stages (Fig. 1f, g). Within the L1 clade, the primate-specific L1PA subfamily displayed the strongest repression, particularly *L1PA2*, which underwent a dramatic 128.90-fold downregulation^3^ (Fig. 1f). Approximately two-thirds of ERV subfamilies followed similar patterns to L1, showing progressive downregulation throughout oogenesis (Fig. 1f). For instance, *HERV-L*, a human specific ERV, exhibited the most substantial repression with an overall 89.43-fold decrease, followed by *HERV-K*, which demonstrated a 29.50-fold reduction from PrF to MII.

Despite the general trend of TE repression, approximately one-third of ERV subfamilies exhibited increasing expression from the PrF to GV stages, with eight subfamilies showing overall upregulation across oocyte development (Fig. 1f). Among these, *HERVS71* exhibited the most notable increase, with a 3.39-fold upregulation. In comparison to robust silencing observed for L1 and most ERVs, Alu and SVA elements underwent relatively moderate repression, with fold changes of 4.71 and 5.36 from PrF to MII stage, respectively. Collectively, these findings indicate a hierarchical repression pattern during human oogenesis, where L1 and the majority of ERVs undergo extensive silencing, while certain ERVs, Alu and SVA elements remain relatively resilient.

### piRNAs as the predominant small non-coding RNAs (sncRNAs) during human oogenesis

Analyses of small RNA composition demonstrated that piRNAs were the most abundant sncRNAs species, accounting for 80.51% of total sncRNAs at the PrF stage and increasing to 84.19% at MII (Fig. 2a). PCA of piRNA expression profiles effectively distinguished PrF and PF oocytes, while GV and MII stages exhibited highly similar expression patterns (Fig. 2b). Among the PIWI genes, *PIWIL3* was the most abundantly expressed, while *PIWIL1* and *PIWIL2* exhibited relatively lower expression levels, and *PIWIL4* was nearly undetectable (Fig. 2c). Based on their length and 3’ end modifications, piRNAs were categorized into two groups, short-piRNAs (17-24 nt, lacking 3’-O-methylation, PIWIL3-associated) and long-piRNAs (25-35 nt, with 3’-O-methylation, PIWIL1- and/or PIWIL2-associated)^39, 41^. The proportion of short-piRNAs increased from 44.88% at PrF to 73.21% at GV, remaining at 71.28% at MII (Fig. 2a), suggesting a predominant role of short-piRNAs in human oocyte compared to long-piRNAs.

**Figure 2.**
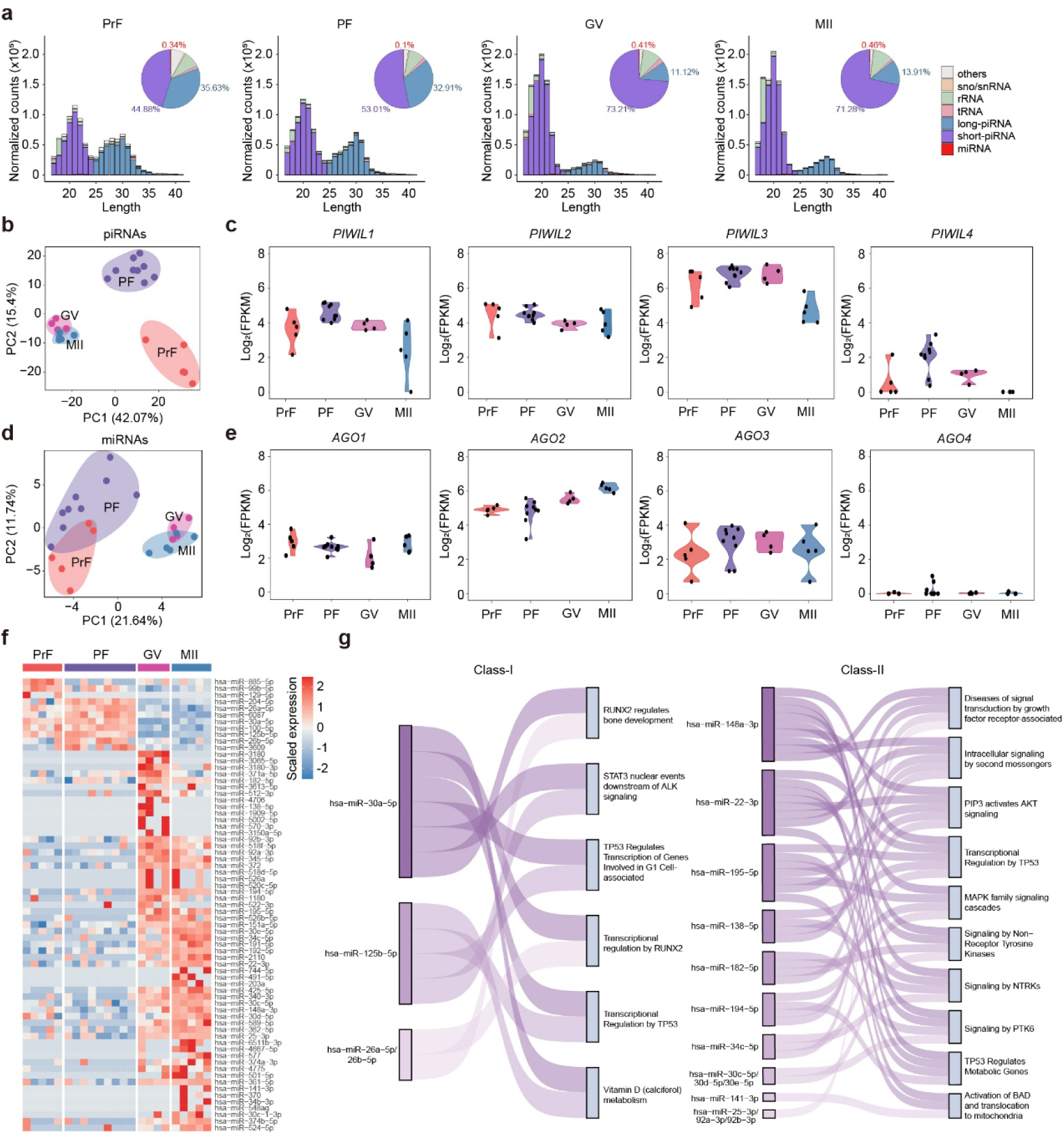
piRNAs are the predominant small non-coding RNAs during human oogenesis. **a.** Distribution of small non-coding RNA categories across four developmental stages of human oocytes. Bar plots represent average proportions from 4-9 biological replicates per stage. Pie charts show the relative contributions of different small non-coding RNA categories at each stage. Expression levels were normalized to reads per million (RPM). **b.** PCA of piRNAs expression in individual oocytes. piRNAs with total expression <100 RPM across all samples were excluded. RPM-normalized expression values were used. **c.** Violin plots showing log_2_-transformed expression levels (FPKM) of PIWI family genes: *PIWIL1, PIWIL2, PIWIL3*, and *PIWIL4,* across oocyte development and maturation. **d.** PCA of miRNA expression in individual oocytes. miRNAs with total expression <100 RPM across all samples were excluded. RPM-normalized expression values were used. **e.** Violin plots showing log_2_-transformed expression levels (FPKM) of AGO family genes: *AGO1, AGO2, AGO3*, and *AGO4,* across oocyte development and maturation. **f.** Heatmap showing stage-specific expression patterns of feature miRNAs enriched at different stages of human oocytes. **g.** Sankey diagrams representing Reactome pathway enrichment of predicted target genes of feature miRNAs, as predicted by TargetScan and miRTarBase. Class I includes miRNAs enriched in PrF and PF stages; Class II includes miRNAs enriched in GV and MII stages.

Unlike in mouse oocytes, endo-siRNAs were undetectable at any sequenced developmental stage of human oocytes, aligning with observations in golden hamsters and other mammals^39, 43-45^ (Fig. 2a). miRNAs, which constituted 0.34% of total sncRNAs in the PrF stage oocytes, exhibited a modest increase after the GV stage (Fig. 2a). miRNA expression profiles grouped the oocytes into two distinct clusters: PrF/PF and GV/MII (Fig. 2d and Supplementary Data 6). Feature analyses identified 11 miRNAs highly enriched in PrF and PF stage oocytes, while 60 miRNAs were upregulated at GV/MII stage oocytes (Fig. 2f). Reactome pathway enrichment analysis revealed that these miRNAs are involved in pathways related to RUNX2, ALK, and TP53, as well as growth factor receptor signaling, second messenger systems, and MAPK signaling (Fig. 2g). Among the ARGONAUTE (AGO) genes, *AGO2* was predominantly expressed, with expression progressively increasing during human oogenesis, while *AGO1* and *AGO3* exhibited consistently low expression (Fig. 2e). *AGO4* was barely detectable in any analyzed stage, in line with its established distinctive role in spermatogenesis^48, 49^. Although miRNAs were expressed at lower levels in oocytes than in somatic cells, their stage-specific expression, the progressive upregulation of AGO2, and enrichment of their targets in critical signaling pathways suggest that miRNAs may contribute to oocyte development and warrant further investigation. Overall, these findings underscore the dominant role of piRNAs, particularly short-piRNAs, in human oogenesis.

### Evolutionarily conserved TE preference of short-piRNAs and long-piRNAs

To explore the potential contribution of piRNAs to global TE repression, we mapped short- and long-piRNAs separately to representative human TE sequences on both sense and antisense strands. Remarkably, approximately 80% of long-piRNAs were associated with ERVs across all developmental stages, compared to around 50% of short-piRNAs (Fig. 3a, b). This preference was largely driven by antisense-aligned long-piRNAs. At the PrF stage, 80.64% of all TE-aligned long-piRNAs were antisense ERV-related, compared to only 5.86% for sense ERV-related, resulting in an antisense-to-sense ratio greater than 13. Although this ratio decreased as development progressed, the bias toward antisense persisted, with a 4.73-fold preference at the MII stage. In contrast, short-piRNAs exhibited a broader association across various TE superfamilies, accompanied by a more balanced antisense-to-sense ratio, generally ranging between 2 to 3 throughout oogenesis (Fig. 3a, b).

**Figure 3.**
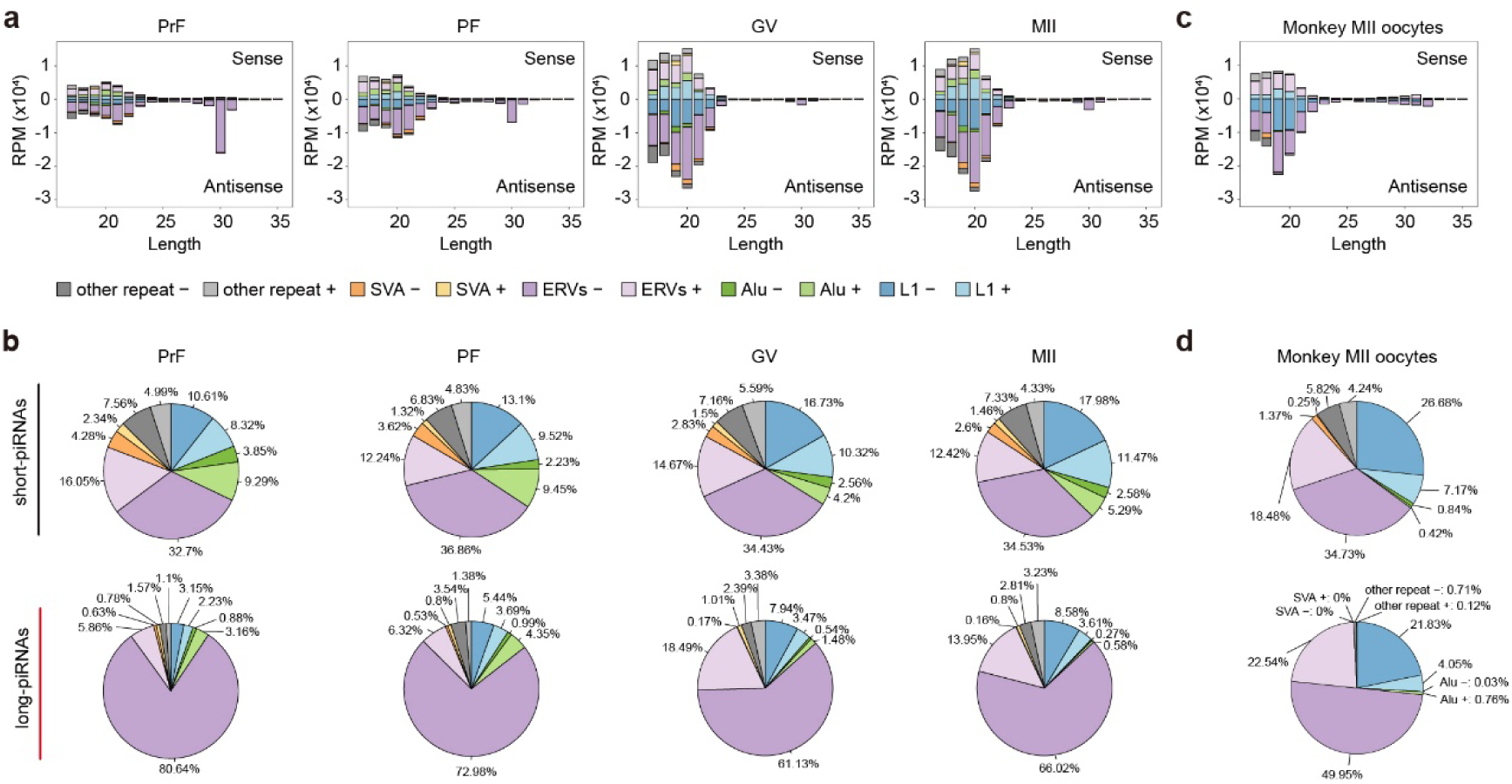
Conserved and biased regulatory roles of short- and long-piRNAs in targeting distinct TE superfamilies during mammalian oogenesis. **a.** Bar plots showing the distribution of piRNAs (17–35 nt) aligned with sense (positive values) and antisense (negative values) strands of major TE superfamilies across human oocyte developmental stages. Each bar represents the average expression (RPM) from 4–9 oocytes per stage. **b.** Pie charts depicting the relative proportions of TE superfamilies targeted by short-piRNAs (17–24 nt, upper panel) and long-piRNAs (25–35 nt, lower panel) at different stages of human oocyte development. Each chart represents the average of 4–9 oocytes, as in (a). **c.** Bar plots showing the distribution of piRNAs (17–35 nt) aligned with sense and antisense strands of major TE superfamilies in monkey oocytes at the MII stage. **d.** Pie charts showing the relative proportions of TE subfamilies aligned with short-piRNAs (17–24 nt, upper panel) and long-piRNAs (25–35 nt, lower panel) in monkey MII oocytes. Each chart represents the average of four biological replicates. Color legends in (b), (c), and (d) are consistent with those used in (a).

Notably, the preference for ERV association among long-piRNAs was conserved across species. In monkey and golden hamster MII oocytes, 72.49% and 69.69% of long-piRNAs, respectively, were ERV-related, compared to 53.21% and 43.87% of short-piRNAs (Fig. 3c, d and Supplementary Fig. 2a, b). In addition, the antisense-to-sense strand ratio for ERV-related long-piRNAs exceeded 10 in golden hamster oocytes, whereas there was almost no bias for ERV-related short-piRNAs (Supplementary Fig. 2a, b). Collectively, these results suggest that long-piRNAs exhibit a conserved, preferential targeting of ERVs across species, indicating their specialized role in ERV repression. In contrast, short-piRNAs exhibit broader TE targeting, likely reflecting their more generalized silencing function across multiple TE families.

### The widespread role of short-piRNAs in TE repression

Analyses of TE-aligned short-piRNA expression revealed three distinct groups corresponding to the PrF, PF, and GV/MII stages (Fig. 4a and Supplementary Data 4). A marked upregulation of short-piRNA was observed from PrF to GV stages during oogenesis (Fig. 4b, c). From PrF to PF, 162 TE subfamilies exhibited significant increases in short-piRNAs expression, nearly twice as many as those showing downregulation. From PF to GV, 130 TE subfamilies showed upregulated short-piRNAs, while only seven were downregulated (Fig. 4c and Supplementary Data 7). Notably, the upregulation of short-piRNAs preceded the onset of substantial TE suppression, which was primarily initiated after the PF stage (Fig. 1d, e), indicating a temporal delay in the repressive effects of short-piRNAs on TE activity at the mRNA levels.

**Figure 4.**
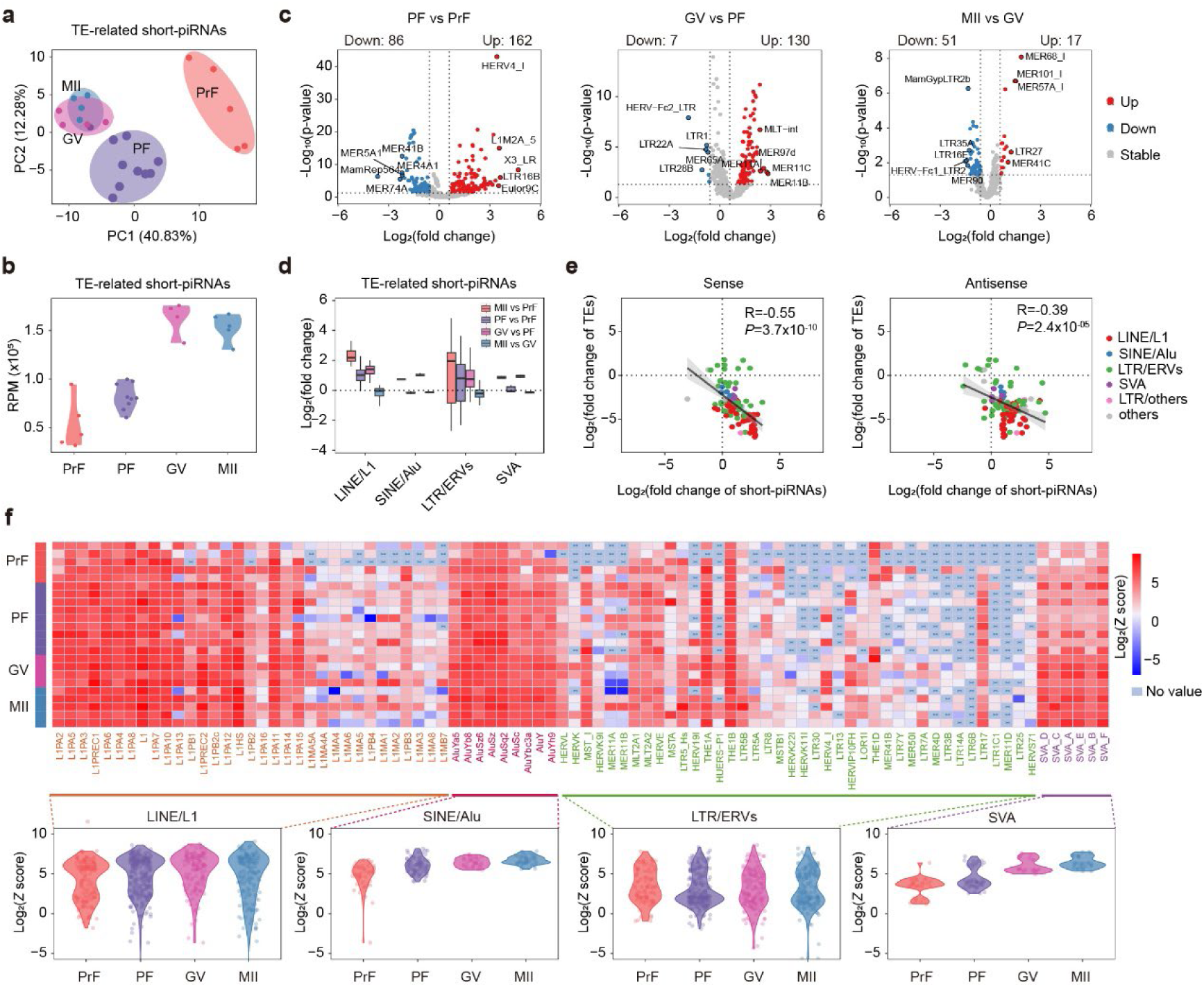
Global upregulation of TE-aligned short-piRNAs contributes to TE repression during human oogenesis. **a,** PCA of TE-aligned short-piRNAs expression across individual human oocytes at different developmental stages. Expression levels were normalized to RPM. **b,** Total abundance (RPM) of TE-aligned short-piRNAs progressively increases throughout oocyte development. **c,** Differential expression analysis of TE-aligned short-piRNAs across adjacent developmental stages (PF vs PrF, GV vs PF, MII vs GV). Significantly up- and downregulated TE-aligned short-piRNAs were identified as in Figure 1e. The number of TE subfamilies with significantly up- and downregulated short-piRNAs, as well as the top five with the highest fold changes, are indicated. **d,** Box plots showing overall fold changes of short-piRNAs aligned to different TE superfamilies across oocyte development from PrF to MII stages, as well as comparisons between adjacent stages (PF vs PrF, GV vs PF, MII vs GV). All TE subfamilies with significantly up- or downregulated short-piRNAs, and all SVA subfamilies regardless of statistical significance, are included. **e,** Scatter plot showing log_2_-transformed fold changes in mRNA levels of differentially expressed TE subfamilies and their aligned short-piRNAs (sense: left; antisense: right) from PrF to MII stages. Points are color-coded by TE superfamily. Pearson correlation coefficients and p-values are shown. **f,** *Upper panel*: Heatmap showing Ping-Pong signature for TE subfamily-aligned short-piRNAs. The Ping-Pong signature was defined by the Z score of 10-nt overlaps between piRNAs on opposite strands. Z score > 1.96 corresponds to p-value < 0.05. Only TE subfamilies with significantly altered mRNA expression from PrF to MII stages were included. *Lower panel*: Violin plots summarizing the changes in Ping-Pong signature scores across oocyte development for each TE superfamily.

Among TE superfamilies, L1-related short-piRNAs exhibited the most pronounced upregulation throughout oogenesis, aligning with L1 being the most substantially downregulated TE superfamily (Fig. 4d, Fig. 1g, and Supplementary Fig. 3a). Following L1, ERV-related short-piRNAs displayed the second most prominent upregulation, although considerable heterogeneity was detected across different ERV subfamilies, mirroring the varied repression patterns observed in these elements. In addition, Alu- and SVA-related short-piRNAs showed relatively modest increases, mainly during the PF-to-GV transition, consistent with their comparatively moderate repression (Fig. 4d and Supplementary Fig. 3a). A robust inverse correlation between TE repression levels and the upregulation levels of corresponding short-piRNAs further supports the contribution of short-piRNAs to global TE silencing (Fig. 4e).

The Ping-Pong amplification mechanism is essential for the post-transcriptional TE repression via PIWI/piRNA-mediated cleavage of target RNAs, generating complementary piRNA pairs with a characteristic 10-nt overlap at their 5’ ends^50, 51^. This process also exhibits an uracil (U) preference at the first nucleotide (1U) of antisense piRNAs, and an adenine (A) bias at the tenth nucleotide (10A) in sense. Indeed, high 1U and 10A preferences were observed in retrotransposons-associated piRNAs (sense strand 1U: 89.14%, 10A: 62.85%; antisense strand 1U: 87.86%, 10A: 44.31%, Supplementary Fig. 3b, c). Specifically, short-piRNAs related to most of the L1, Alu, and SVA subfamilies showed strong Ping-Pong signatures (Fig. 4f), with the intensity of these signatures correlating with their progressive repression throughout oocyte development. However, this signature was less prevalent among ERV-related short-piRNAs, with notable exceptions such as THER1A and THER1B. Collectively, these observations suggest that the PIWIL3-mediated Ping-Pong pathway may contribute to the extensive regulatory influence of short-piRNAs on TE repression during human oogenesis.

### Collaborative regulatory role of long-piRNAs in repressing ERVs

TE-aligned long-piRNA expression segregated oocytes into three distinct groups: PrF, PF, and GV/MII stages (Fig. 5a and Supplementary Data 4). However, long-piRNAs were less effective than short-piRNAs in distinguishing PrF from PF stage oocytes. Unlike the widespread upregulation of short-piRNAs during oogenesis, TE-aligned long-piRNAs showed a declining trend in expression (Fig. 5b, c). Differential expression analysis revealed a substantial decrease in TE-aligned long-piRNAs between the PF and GV stages, with 58 TE subfamilies exhibiting significant downregulation of their corresponding long-piRNAs, compared to 13 subfamilies that were upregulated (Fig. 5c and Supplementary Data 8). Despite the overall decline, 18 ERV subfamilies and two derivative LTR elements exhibited significant upregulation of their associated long-piRNAs, aligning with the preferential ERV targeting by long-piRNAs (Fig. 5d and Fig. 3a, b). The Ping-Pong signature was either undetectable or substantially diminished for most TE-aligned long-piRNAs (Supplementary Fig. 4a), including those ERVs or solo LTRs with upregulated long-piRNAs (Supplementary Fig. 4b).

**Figure 5.**
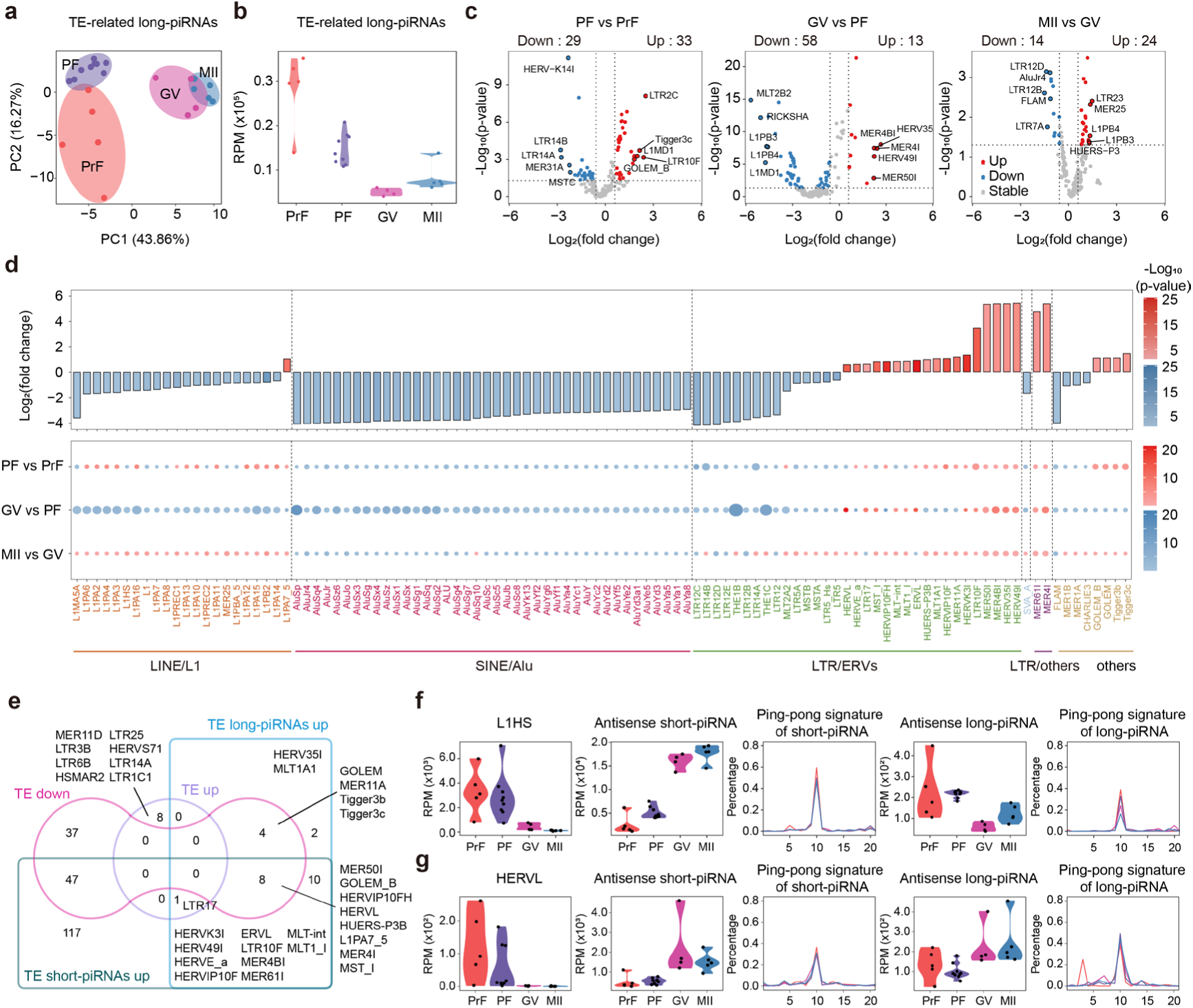
TE-aligned long-piRNAs play an ERV-specific repression role during human oogenesis. **a,** PCA of TE-aligned long-piRNA expression across individual oocytes at different developmental stages. Expression levels were normalized to RPM. **b,** Total abundance (RPM) of TE-aligned long-piRNAs gradually decreases from the PrF to GV stages. **c,** Differential expression analysis of TE-aligned long-piRNAs between adjacent developmental stages (PF vs PrF, GV vs PF, MII vs GV). TE subfamilies with significantly aligned long-piRNAs were identified as in Figure 1e. The number of TE subfamilies with significantly up- and downregulated long-piRNAs, as well as the top five with the highest fold changes, are indicated. **d,** *Upper Panel*: Bar plot showing log_2_-transformed fold changes and corresponding p-values of the TE subfamilies with differentially expressed long-piRNAs between the PrF and MII stages. *Lower Panel*: Fold changes and p-values for these same TE subfamilies across intermediate stages (PF vs Prf, GV vs PF, MII vs GV). Circle size reflects the magnitude of fold change; color intensity reflects -log_10_(p-value). Dashed lines indicate TE subfamilies by superfamily. **e,** Overlap analysis showing shared and distinct TE subfamilies that were up- and down-regulated at the mRNA level or showed upregulation of aligned short-piRNAs or long-piRNAs. The number of TE subfamilies in each intersection group is indicated. **f, g**, Expression dynamics and Ping-Pong signature of representative TE subfamilies *L1HS* (f) and *HERVL* (g) throughout the development of human oocytes. For *L1HS* and *HERVL*: mRNA expression, aligned antisense short-piRNAs expression, Ping-Pong signature of short-piRNAs, aligned antisense long-piRNAs expression, and Ping-Pong signature of long-piRNAs are shown.

To examine the cooperative contributions of short- and long-piRNAs to TE repression during human oogenesis, we assessed the overlap among TEs with significantly altered mRNA levels (upregulated or downregulated), TEs with upregulated short-piRNAs, and those with upregulated long-piRNAs (Fig. 5e). Of the 96 downregulated TE subfamilies, 61.46% (59 out of 96) were associated with either upregulated short- or long-piRNAs. Among these, 93.22% (55 out of 59) were associated with upregulated short-piRNAs, while 20.34% (12 out of 59), most of which belonged to ERV subfamilies, were linked to upregulated long-piRNAs. *L1HS*, *AluYb8*, and *SVA_A* represent TE subfamilies that are primarily regulated by short-piRNAs with repression correlated inversely with a gradual increase in corresponding antisense short-piRNAs and accompanied by prominent Ping-Pong signatures (Fig. 5f and Supplementary Fig. 4b, c). Among the TE subfamilies associated with long-piRNAs, 66.67% (8 out of 12) also showed upregulation of short-piRNAs, suggesting a coordinated role for both piRNA classes in repressing these elements. *HERV-L*, one of the most strongly downregulated ERV subfamilies, showed an inverse correlation with both antisense short- and long-piRNAs during oocyte development (Fig. 5g). This repression was accompanied by a weak but detectable Ping-Pong signature. All these observations support a dominant role for short-piRNAs in broad TE silencing, with long-piRNAs contributing to ERV-specific repression.

In addition, we identified eight TE families, including *HERVS71, LTR25, MER11D, LTR1C1, LTR17, LTR6B, LTR14A,* and *LTR3B,* that remained transcriptionally active during human oogenesis (Fig. 1f and Fig. 5e). Neither short-piRNAs nor long-piRNAs associated with these elements showed significant upregulation. Notably, seven of these elements, with the exception of *LTR1C1,* are specific to the human genome. The limited or absent repression of these evolutionarily younger elements by the piRNA/PIWI may reflect a lag in the adaptation of the piRNA/PIWI system to those recently emerged transposon threats.

### Highly productive piRNA clusters evolved asymmetric antisense bias toward L1 and ERVs

Primary piRNA biogenesis is tightly associated with specific genomic loci across various species^28, 50, 52^. To determine the genomic origins of piRNAs expressed during human oogenesis, we employed a proximity-based clustering strategy adapted from the piClust algorithm^53^. Using a threshold of at least four uniquely mapped piRNAs within a 2.5-kilobase (kb) window and merging overlapping regions, we identified 14,603 piRNA-generating clusters. Remarkably, the top 20 most highly expressed clusters accounted for 91.72 % of all uniquely mapped piRNAs and exhibited stage-specific expression dynamics (Fig. 6a, b, and Supplementary Data 9). These clusters generated both short- and long-piRNAs, with short-piRNA production generally increasing as oogenesis progressed (Fig. 6c).

**Figure 6.**
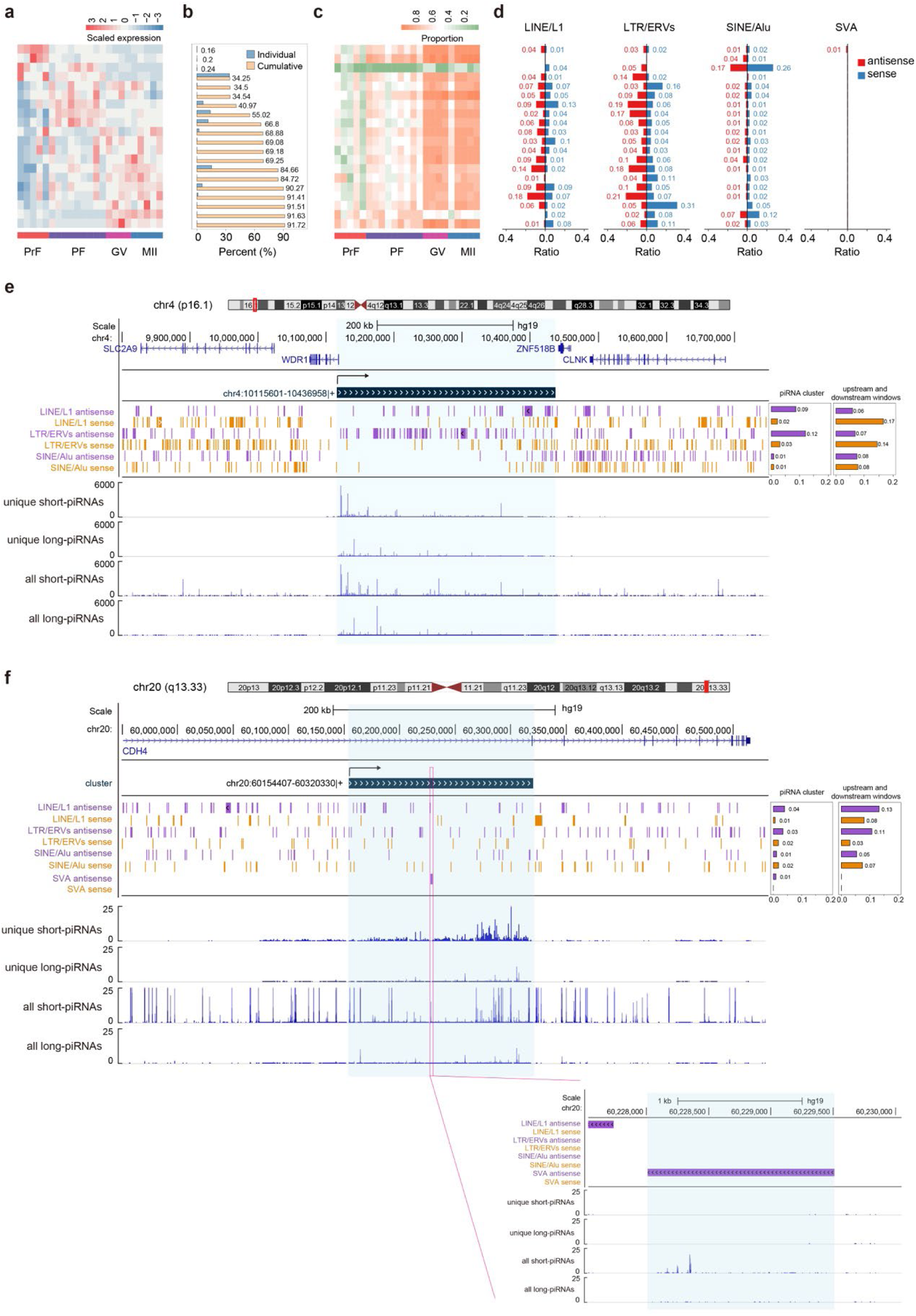
Highly productive piRNA clusters exhibit asymmetric enrichment for TE families, particularly antisense L1s and ERVs. **a,** Heatmap displaying expression profiles of the top 20 most highly expressed piRNA clusters, which collectively account for 92.50% of uniquely mapped piRNAs across all sequenced human oocytes. Expression values were row-scaled. **b,** Bar plot showing the proportion of piRNAs generated by each cluster (from panel a) and the cumulative contribution of these clusters to the total uniquely mapped piRNA pool. **c,** Percentage of short-piRNAs relative to the total uniquely mapped piRNAs produced by each of the top 20 clusters (from panel a). **d.** Strand-specific bar plots illustrating the insertion frequencies of LINE/L1, SINE/Alu, LTR/ERV, and SVA elements within the top 20 clusters (from panel a). **e.** Genome browser view of the most highly expressed piRNA cluster (chr4:10115601-10436958, + strand), displaying surrounding gene annotations, TE superfamily density, and both uniquely and total mapped piRNA expression levels (short-piRNAs and long-piRNAs). **f.** Genome browser view of another top piRNA cluster with an antisense SVA insertion (chr20:60154407-60320330, + strand), with the same annotations and tracks as in (e). In (e) and (f), TE annotations were obtained from RepeatMasker, and piRNA expression was normalized to RPM. Right-side bar plots summarize the orientation (sense vs. antisense) of TE superfamily insertions within each cluster and their upstream/downstream flanking windows.

TE enrichment analysis revealed a strong bias toward L1 and ERV fragments within the top 20 clusters, particularly for the antisense strand, when compared to genomic background regions (Fig. 6d and Supplementary Fig. 5 and Supplementary Data 10). This asymmetric insertion pattern facilitated the preferential generation of antisense-aligned piRNAs targeting L1 and ERVs (Supplementary Fig. 5a, b). This phenomenon is reminiscent of the *flamenco/COM* locus in *Drosophila*, which generates antisense piRNAs to silence *gypsy, Idefix,* and *ZAM* elements^50, 54, 55^. In contrast, Alu elements were not enriched in these top clusters compared to genomic background regions (Fig. 6d and Supplementary Fig. 5c), consistent with the relatively low production of Alu-derived piRNAs and their modest repression during oocyte development.

The most productive piRNA cluster, spanning approximately 320 kb between *WDR1* and *ZFN518B* on chromosome 4, contributed 34.05% of all uniquely mapped piRNAs. This unidirectionally transcribed cluster contained abundant antisense insertions of L1 and ERVs (Fig. 6e) and generated numerous short- and long-piRNAs targeting these elements (Supplementary Fig. 5). Similarly, the second most productive cluster, also on chromosome 4 (∼300 kb between *CCNG2* and *CXCL13*), contributed 15.41% of uniquely mapped piRNAs and displayed a comparable enrichment for antisense insertion of L1 and ERVs (Supplementary Fig. 6a). By contrast, the third most productive cluster featured minimal L1 context but was enriched for ERVs, producing predominantly antisense piRNAs targeting ERVs (Supplementary Fig. 6b). This suggests that this cluster primarily functions as a specialized regulator of ERVs, distinct from the broader TE repression roles observed in the top two clusters.

SVA, a relatively young TE element with approximately 4,000 annotated copies in the human genome, was represented by antisense insertions in one of the top 20 clusters. This insertion was accompanied by the production of SVA-targeting antisense piRNAs (Fig. 6d, f), suggesting that this cluster may serve as a SVA suppressor. Conversely, L2 elements, which are largely inactive in the human genome, were absent from the top clusters. These patterns suggest that evolutionary pressures preferentially shape the composition and activity of piRNA clusters based on the functional relevance and activity of specific TE families (Supplementary Fig. 6). Taken together, these findings underscore the unique evolutionary dynamics of human piRNA clusters and their adaptive specialization in silencing diverse and evolving TE populations during oocyte development.

## Discussion

To maintain genomic integrity, the host genome has evolved a range of transcriptional and post-transcriptional defense systems to suppress TE activity, such as the piRNA pathway, APOBEC3 family, and Kruppel-associated box (KRAB) zinc finger gene families^56, 57^. In this study, we simultaneously profiled the small and long RNA transcriptomes of human oocytes across four key developmental stages, providing a comprehensive view of the distinct roles of short- and long-piRNAs in TE silencing. Our findings emphasize the broad and potent suppressive effects of piRNAs on TEs during human oogenesis, with over 60% of downregulated TEs correlated with the upregulation of short- and/or long-piRNAs.

Among TE superfamilies, L1 was the primary target of piRNAs and exhibited the most pronounced repression throughout oocyte development. This effect was particularly evident in L1PA subfamilies, which remain active in the human genome and continually result in insertional polymorphisms in human populations^58, 59^. Interestingly, Alu and SVA elements, which depend on L1 activity for mobilization, displayed only modest overall repression. This finding suggests that while L1 elements are effectively targeted by piRNAs, the suppression of non-autonomous TEs like Alu and SVA may rely more heavily on L1 regulation rather than direct piRNA targeting.

Although ERVs have long been considered largely inactive in humans due to their limited transpositional capacity, recently emerging evidence suggests that their reactivation may contribute to pathological conditions, including cancers, autoimmune diseases, and neurodegenerative disorders^60-65^. Among the various ERV families, *HERV-K* is particularly notable for exhibiting signs of recent activity and contributing to individual genomic variation^8, 66^. Its transcriptional activity and potential for protein production have been observed in several pathological contexts^67, 68^, highlighting its relevance to human disease. In this study, we observed robust silencing of ERVs, particularly *HERV-L* and *HERV-K*. Both short- and long-piRNAs contributed to ERVs repression, with long-piRNAs showing a stronger preference for ERVs targeting. These observations align with functional studies in golden hamsters, where long-piRNAs play a more pivotal role in ERV repression than short-piRNAs. Depletion of PIWIL1-associated long-piRNAs in golden hamsters leads to ERV accumulation and sterility, while PIWIL3/short-piRNAs can compensate for L1 suppression but not ERV regulation^43^. Conversely, depletion of PIWIL3/short-piRNAs results in subfertility without aberrant ERV activation, likely due to the compensatory function of long-piRNAs in ERV silencing^46^.

The differential activity of TEs across species may underlie the functional divergence of the PIWI/piRNA pathways in mammalian oogenesis. In rodents, including golden hamsters, ERVs are the most active autonomous TEs, whereas in humans, L1 elements are the dominant autonomous transposons^45, 69^. This evolutionary shift may also explain why short-piRNAs play a predominant role in suppressing L1 activity during human oogenesis, potentially reducing the reliance on long-piRNAs in this context. Consequently, disruptions in short-piRNA biogenesis are hypothesized to have more severe consequences for female fertility than long-piRNA deficiencies. Further clinical and genetic studies are required to fully elucidate the functional impact of PIWI/piRNAs disruptions on human reproduction.

## Materials and Methods

### Ethics statement

The study was approved by the Reproductive Study Ethics Committee of Shanghai Zhongshan Hospital (approval number: B2018-244). Written informed consent was obtained from all participants prior to oocyte donation, following clinical protocols of the Department of Assisted Reproduction at Zhongshan Hospital, affiliated with Fudan University, Shanghai, China.

### Human oocyte collection

Human follicles were isolated from fresh ovarian cortex obtained from patients undergoing ovarian cyst or cystic teratoma resection, following previously described procedures^70^. Briefly, ovarian cortical tissues were fragmented into pieces measuring 30–600 μm and digested with 1 mg/mL collagenase type IA (Sigma-Aldrich) and 8 IU/mL DNase I (Sigma-Aldrich) in 10 mL of M199 medium (Gibco) at 37°C for 30 minutes with gentle agitation. Follicles were subsequently isolated and evaluated under a stereomicroscope (Nikon SMZ 1000), and their diameters were measured using an inverted microscope (Nikon Ti2).

Ovarian stimulation and oocyte collection were performed as previously described^41, 70^. GV oocytes were collected following stimulation. MII stage oocytes were obtained after overnight incubation to facilitate maturation, characterized by the presence of a distinct first polar body under microscope observation. Granulosa cells were removed via incubation with 1 mg/mL hyaluronidase. The zona pellucida was eliminated using Acidic Tyrode’s Solution (Millipore), yielding denuded oocytes, which were then washed and transferred to PCR tubes containing 4 μL of cell lysis buffer (0.2% Triton X-100, Sigma; 1 U/μL RiboLock RNase inhibitor, Thermo Fisher). The oocytes were stored at -80°C until further processing.

### Single-oocyte small-RNA library construction

Single-oocyte small-RNA libraries were constructed as previously described^71^. Briefly, lysates from the single oocyte were incubated at 72 °C for 3 minutes to release and unfold the small RNAs. After 3′ adapter ligation, 5 U of lambda exonuclease and 25 U of 5′ deadenylates were used to remove the excess 3′ adapter. Subsequently, the 5′ adapter was ligated, followed by reverse transcription and pre-amplification. For final amplification, 1 μL of the pre-amplified product was used as a template. Libraries were size-selected by electrophoresis on a 6% polyacrylamide gel, and 130–160 bp DNA fragments were extracted to recover small RNA libraries. Sequencing was performed on the HiSeq X Ten platform (Illumina) using 2 × 150 bp reads.

### Single-cell mRNA library construction

Single-oocyte mRNA was captured and amplified using Smart-seq2, as previously described^71, 72^. 2 ng of purified cDNA was subjected to a tagmentation reaction with Tn5 transposase^73^. Sequencing was carried out on the Illumina HiSeq X Ten platform with 2x150 bp reads.

### Data processing of small RNA-seq and long RNA-seq

For small RNA-seq, raw read1 sequences were processed using the fastx_toolkit (v.0.0.14) for quality filtering and adapter trimming. Reads lacking the 3’ adapter sequence or shorter than 17 bp post-trimming were discarded. Reads of 17-40 bp were mapped to the human genome (hg19) using Bowtie (v.1.2) with parameters --k=100 -- v=0.

For long RNA-seq, raw pair-end reads were processed using TrimGalore (v.0.5.0) for adapter trimming and quality filtering. The remaining reads were mapped to the human genome (hg19) and representative TE sequences using STAR (v2.7.1a) with parameters --winAnchorMultimapNmax 10000 --outFilterMultimapNmax 10000. Gene abundance was quantified using RSEM (v.1.2.31) with default settings.

### Categorization of small non-coding RNAs

Mapped small RNA-seq reads were further aligned to annotations of small non-coding RNAs in the following order: pre-miRNAs, tRNAs, rRNAs, snoRNAs, and snRNAs, using Bowtie (v.1.2) with parameters --a --v=0 --norc. Reads that precisely matched the 5’ start site of miRNAs and 3’ ends with less than 3 bp deletions or additions derived from pre-miRNAs were considered for miRNA quantification.

Unassigned sequences that were not categorized as pre-miRNA, tRNA, rRNA, snoRNA, or snRNA, with lengths ranging from 17–40 nt, were identified as piRNA clusters using a modified piClust with MinReads=4 and Eps=2500 bp. Overlapping clusters were merged using mergeBed from bedtools (v.2.30.0). Reads mapping within these clusters were defined as piRNA candidates.

piRNA candidates were subsequently aligned to representative TE sequences using Bowtie with the parameters --a --v = 2. Based on alignment orientation, piRNAs were classified as TE-aligned antisense piRNAs (mapping to the opposite strand of TEs) or TE-aligned sense piRNAs (mapping to the same strand of TEs). Reads were further categorized by TE superfamily in the following order: LINE/L1, SINE/Alu, LTR/ERVs, LTR/others, SVA, and others. If a read mapped to multiple TE subfamilies within a superfamily, it was counted simultaneously for each subfamily.

### Cell Clustering and Feature Gene Analysis

Cell clustering based on gene, TE, miRNA, and piRNA expression levels was performed using principal component analysis in Seurat (v.2.3.4). Feature genes for each stage were identified using FindAllMarkers from Seurat, and enriched pathways were analyzed using Gene Ontology website.

### Differential Expression Analysis

Differential expression of TEs and their aligned piRNAs was analyzed using DESeq2, with thresholds of fold change ≥ 1.50 or ≤0.67 and p-value ≤0.05. TEs or aligned piRNAs with average expression levels below 100 RPM across all samples were excluded from this analysis.

### Identification of Ping-Pong signature

Ping-Pong signatures were detected by calculating the Z score of 10-nucleotide overlaps of piRNAs from opposite strands. A Z score > 1.96 was considered significant^74^ (p-value < 0.05).

### Statistics and Reproducibility

No formal statistical method was used to predetermine the sample size. Sample sizes were determined based on prior studies to ensure reliable measurement of experimental parameters according to field standards^43, 75^. Samples were randomly allocated, and blinding was implemented during library construction. Data analyses were conducted using R version 3.6.3. Results are presented as mean ± s.e.m., with statistical tests and p-values provided in figure legends.

### Sources of Annotations

Human (hg19) and monkey (macFas5) genome sequences were obtained from the UCSC Genome Browser, while the golden hamster genome (MesAur1.0) was retrieved from Ensembl. For human and monkey, the representative sequences of TEs were retrieved from RepBase, and genomic TE annotations were sourced from UCSC. For the hamster, TE sequences and annotations were obtained following previously published protocols^43, 75^. Gene annotations for hg19 were sourced from GENCODE (version 19). Functional RNA annotations for human were obtained from miRBase (pre-miRNAs and miRNAs), Genomic tRNA Database (tRNAs), NCBI GenBank (18S and 28S rRNAs), and Ensembl (5S and 5.8S rRNAs, snoRNA and snRNAs).

## Data Accessibility

The sequencing data generated in this study have been deposited in the NCBI GEO database under accession number GSE296363. The data are available for review using the token ojepagsydbsrpgt. Publicly available datasets used in this study include small-RNA sequencing data for golden hamsters from GSE169528^43^ and monkey GV oocytes from GSE95218^41^. Additional processed data related to this study are available from the corresponding author upon request.

## Acknowledgement

The authors are very grateful to all the oocyte donors. We thank Qi Qiu, Zhen Miao, Zhenying Tian (University of Pennsylvania), and Daosheng Phillip Xiao for their insightful comments on the manuscript, and the members of our laboratories for valuable feedback on data analyses.

## Contributions

L.W. and H.Z. conceived the study and secured fundings. M.L. and A.R. were responsible for oocyte collection under the supervision of S.L. H.Z. constructed the small and long RNA-seq libraries. F.Z. and Y.X. conducted bioinformatic analyses and prepared figures. F.Z., H.Z., and L.W. interpreted the data and wrote the manuscript. All authors read and approved the manuscript.

## Supplementary Figures

**Supplementary Figure 1.**
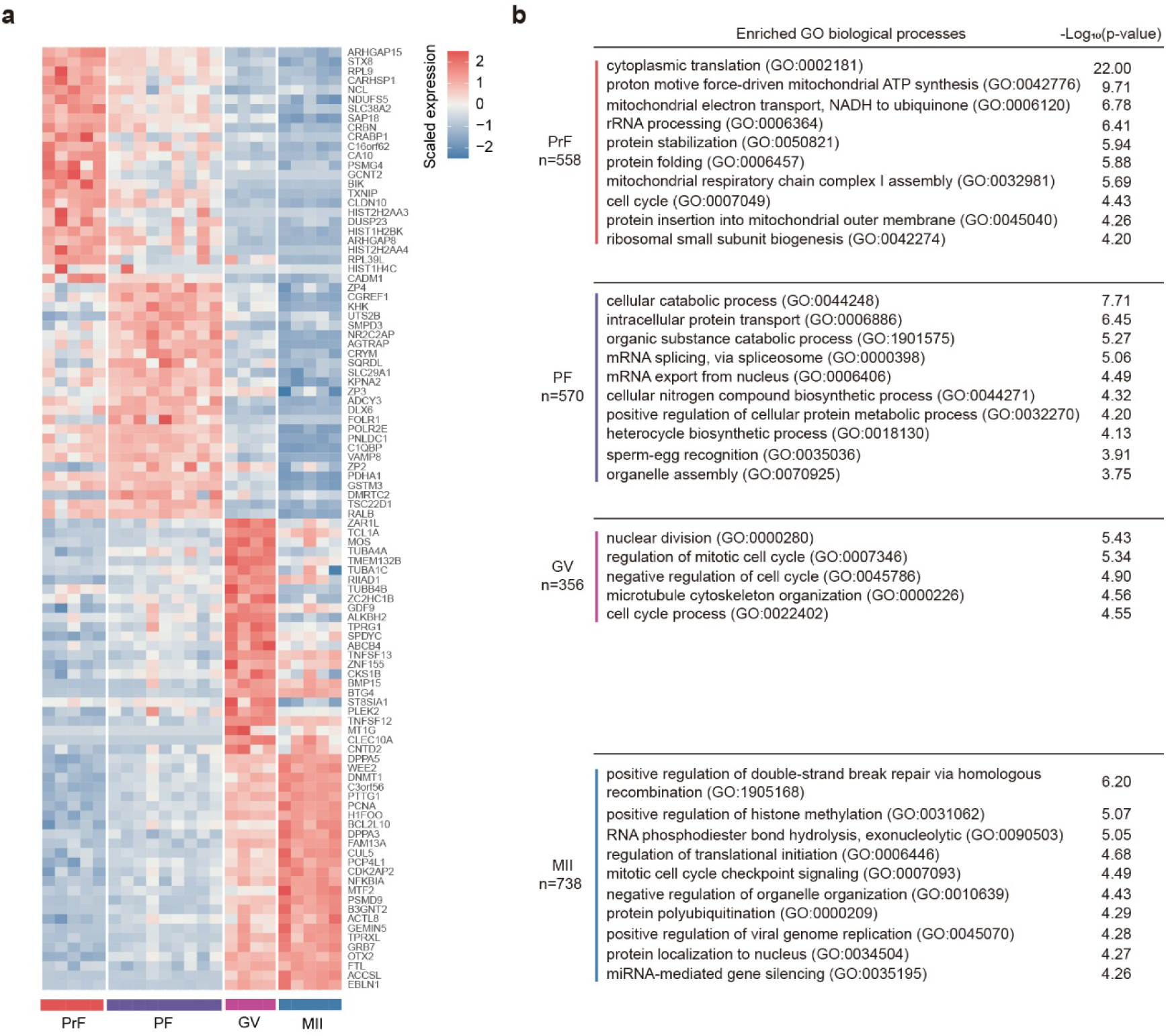
Stage-specific feature genes and enriched biological pathways revealed by long transcriptome profiling of developing human oocytes. **a.** Heatmap displaying the top 25 stage-enriched feature genes across four developmental stages of human oocytes. Gene expression levels were normalized to FPKM and row-scaled. **b.** Top 10 significantly enriched GO biological process items associated with the feature genes at each developmental stage. The number of feature genes identified per stage is indicated. Only five GO items were significantly enriched for the feature genes at the GV stage.

**Supplementary Figure 2.**
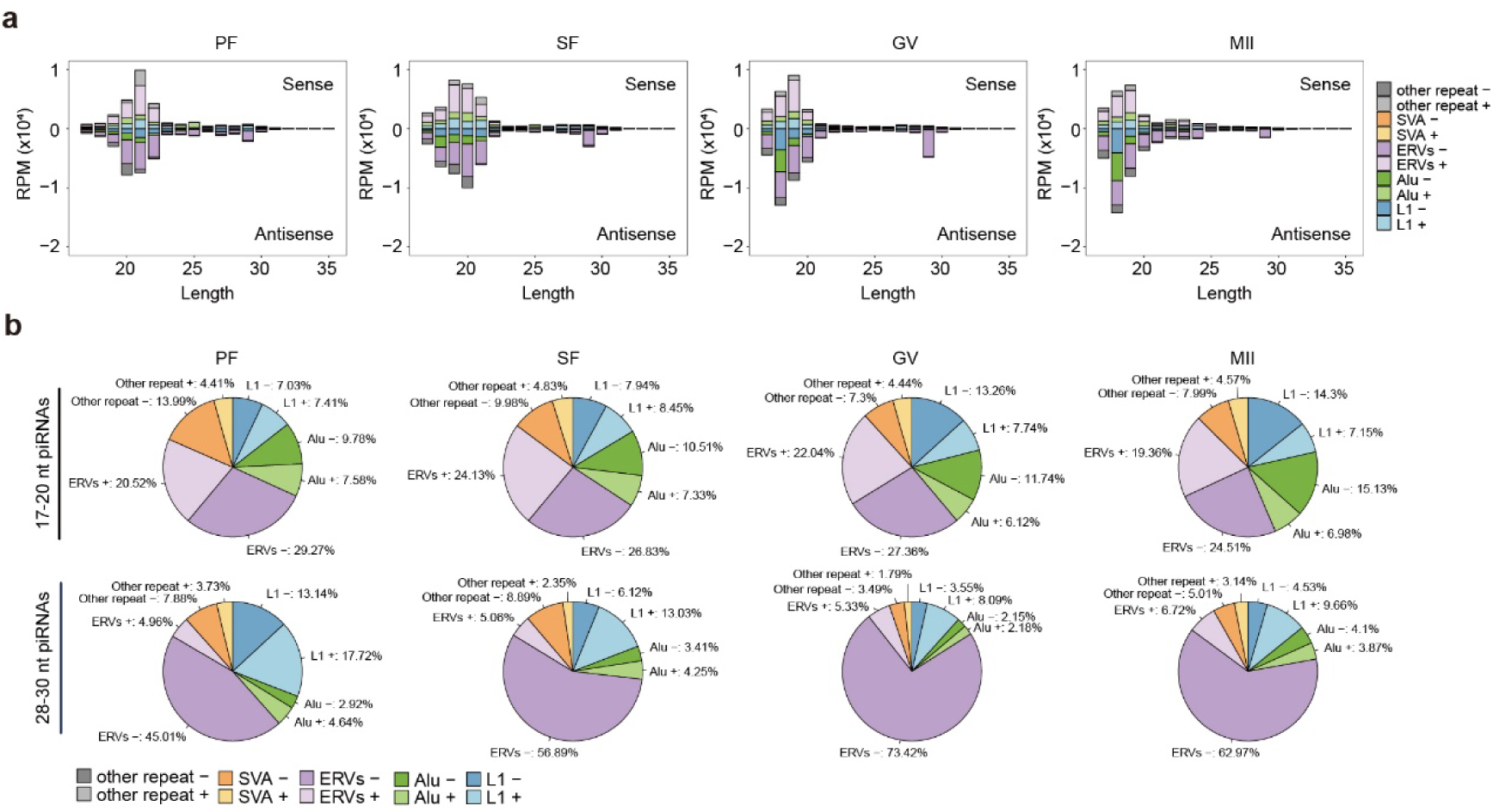
Conserved antisense targeting preference of long-piRNAs for ERV elements in golden hamster oocytes. **a.** Bar plots showing the distribution of piRNAs aligned to sense (positive values) and antisense (negative values) strands of various TE superfamilies in developing, full-grown and mature oocytes of golden hamsters. Each bar represents the average expression (RPM) from 2-4 oocytes per stage. **b.** Pie charts displaying the proportion of TE superfamilies targeted by short-piRNAs (18–20 nt, upper panel) and long-piRNAs (28–30 nt, lower panel) at each developmental stage. Each chart represents the average of 2–4 oocytes per stage, as in (a).

**Supplementary Figure 3.**
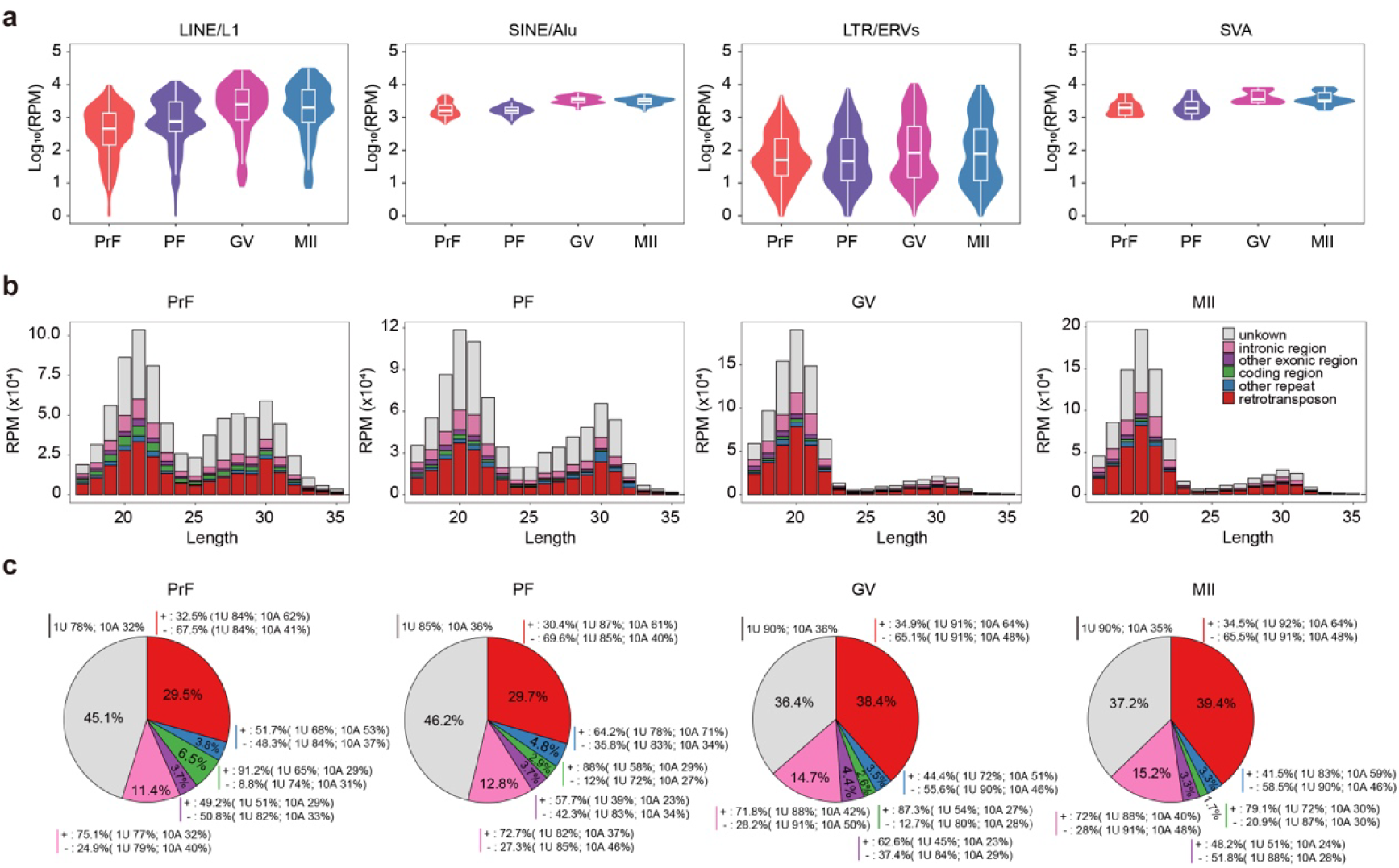
Differential upregulation patterns of TE superfamily-aligned short-piRNAs and presence of Ping-Pong signatures during human oogenesis. **a,** Violin and box plots showing the dynamic upregulation of TE subfamily-aligned short-piRNAs during oocyte development from the PrF to MII stages. **b,** Bar plots illustrating the length distribution of piRNA originating from different genomic regions. **c,** Pie charts showing the proportions of piRNAs from each genomic category in (b), along with the strand-specific 1U and 10A nucleotide biases indicative of the Ping-Pong mechanism.

**Supplementary Figure 4.**
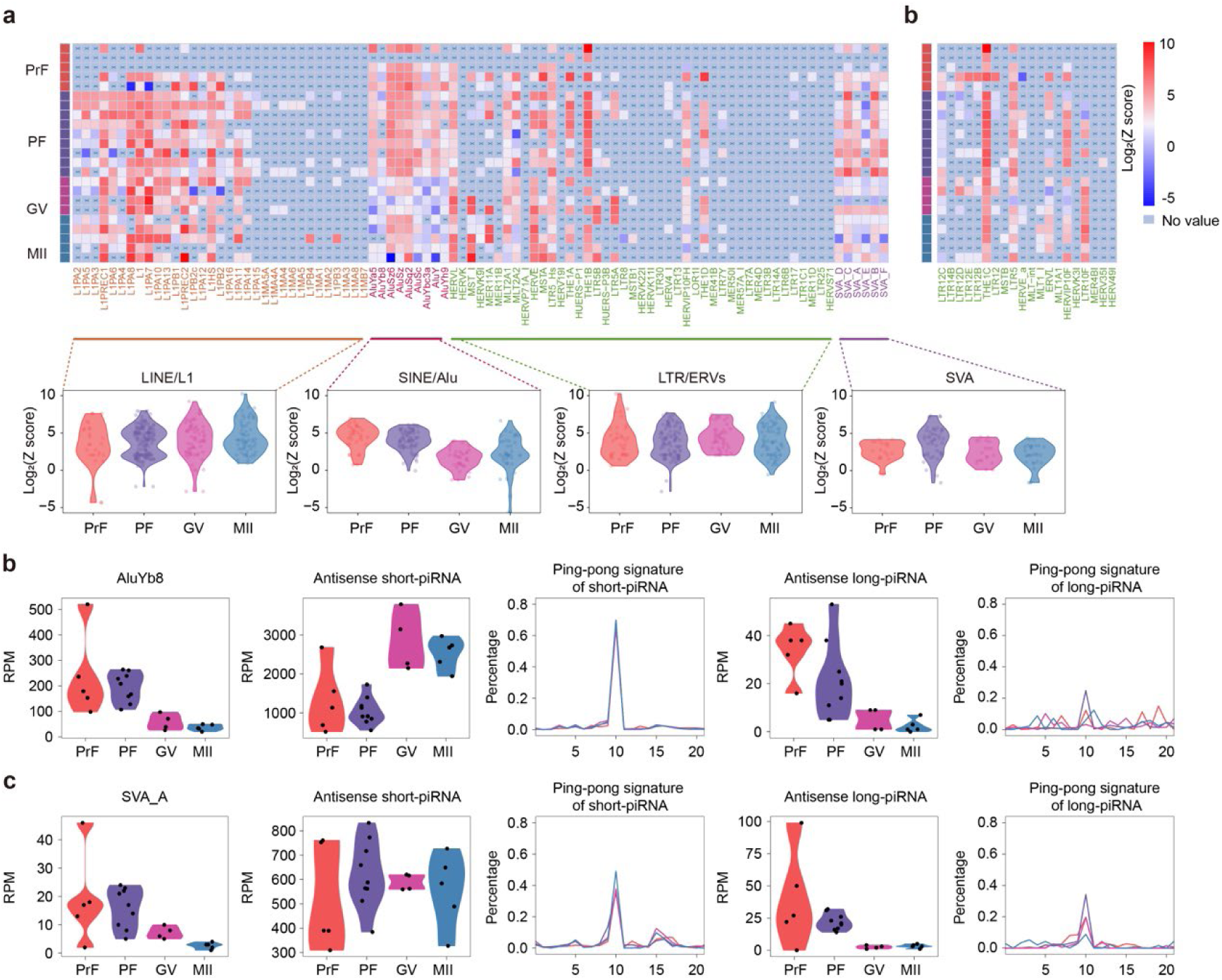
Ping-Pong signature analysis of TE-aligned long-piRNAs and showcasing of Alu and SVA elements regulation patterns during human oogenesis. **a**, *Upper panel*: Heatmap showing Ping-Pong signature scores of TE subfamilies aligned with sense and antisense long-piRNAs. Only TE subfamilies with significantly altered mRNA expression (from Figure 1f) are included. *Lower panel*: Violin plots summarizing Ping-Pong scores across TE superfamilies. **b**, Heatmap showing Ping-Pong signature scores of TE subfamilies with significantly upregulated long-piRNAs identified in Figure 5d. In both (a) and (b), the Ping-Pong signature is assessed using the Z score of 10-nt overlaps between piRNAs on opposite strands (Z > 1.96 corresponds to p < 0.05). **c–d**, Expression dynamics and Ping-Pong signature analysis of representative non-autonomous TE subfamilies AluYa8 (c) and SVA_A (d) during oocyte development. For AluYa8 and SVA_A, the following features are shown: mRNA expression, aligned antisense short-piRNAs expression, Ping-Pong signature of short-piRNAs, aligned antisense long-piRNAs expression, and Ping-Pong signature of long-piRNAs.

**Supplementary Figure 5.**
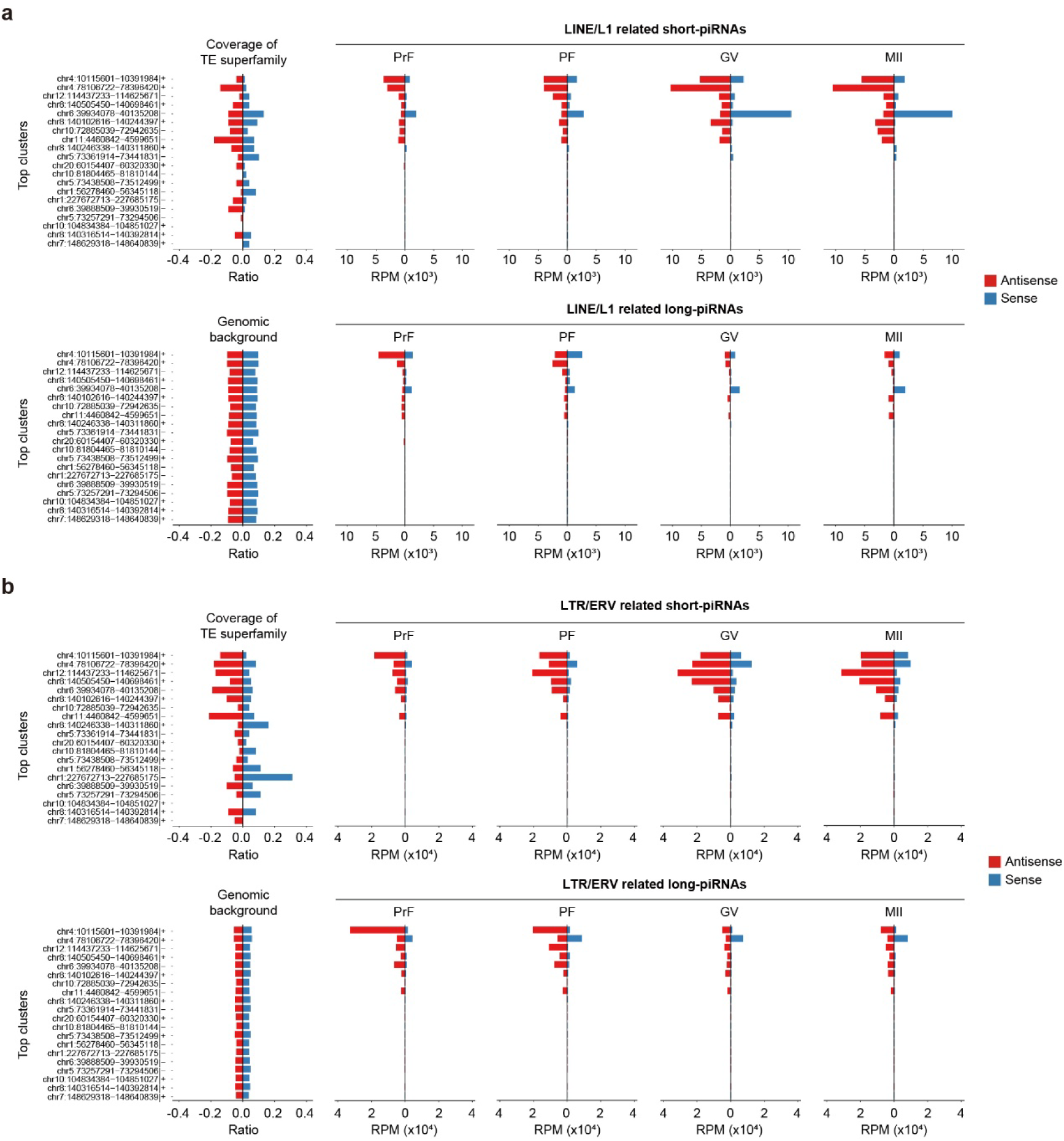
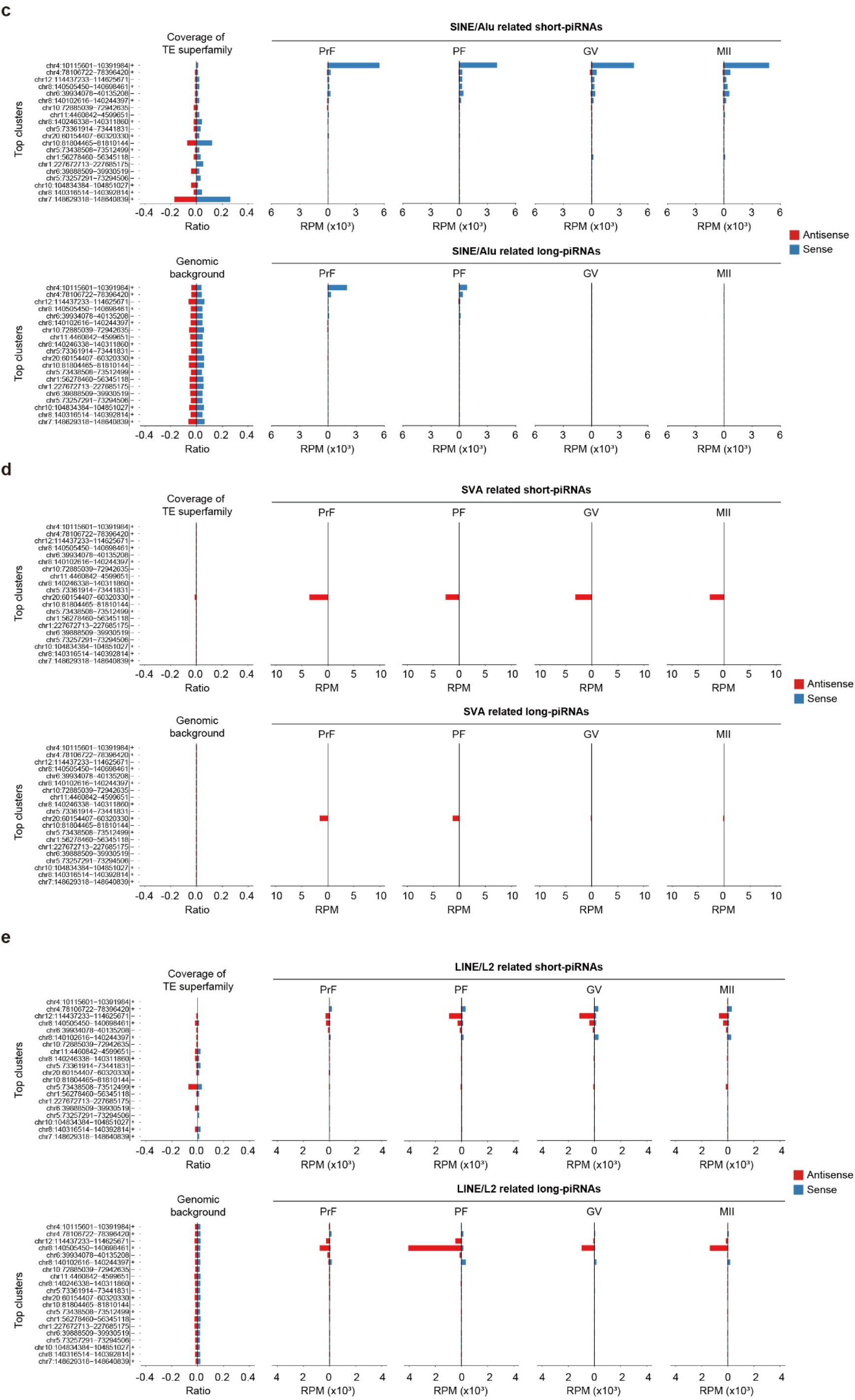
Comparative analysis of TE superfamily enrichment in top 20 piRNA-producing clusters and their aligned short- and long-piRNA expression during human oogenesis. **a–e**, Strand-specific bar plots showing the genomic coverage of LINE/L1 (a), LTR/ERV (b), SINE/Alu (c), SVA (d), and LINE/L2 (e) superfamilies within the top 20 most highly expressed piRNA-generating clusters. Corresponding bar plots show the total abundance (RPM) of aligned sense (blue) and antisense (red) short-piRNAs and long-piRNAs across different oocyte developmental stages. Genomic background coverage was calculated as the average of 1,000 randomized genomic regions of matched length for each cluster.

**Supplementary Figure 6.**
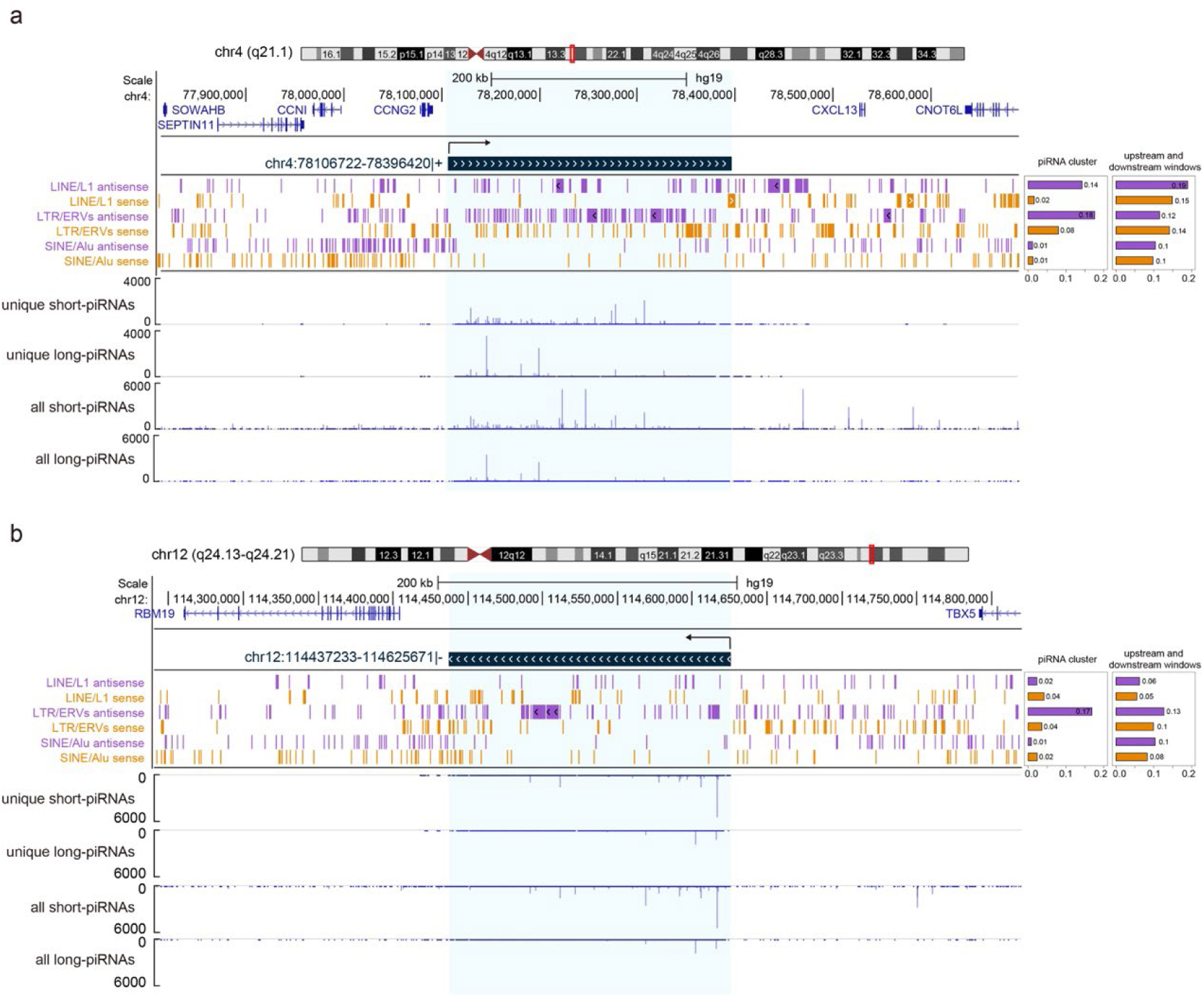
Genomic landscape of top piRNA clusters in human oocytes. **a**, Genome browser view of the second most highly expressed piRNA cluster (chr4:78,106,722–78,396,420: + strand), showing the surrounding gene annotations, density of TE superfamily insertions, and expression levels of uniquely and total mapped short- and long-piRNAs. **b**, Genome browser view of the third most highly expressed piRNA cluster (chr12:114,437,233–114,625,671: – strand), with the same features displayed as in (a). In (a) and (b), TE annotations were obtained from RepeatMasker, and piRNA expression was normalized to RPM. The bar plots on the right summarize the proportion of TE superfamilies inserted in the cluster regions (sense vs. antisense) and their flanking upstream and downstream windows.

## Supplementary Data

**Supplementary Data 1**

Summary of sample information.

**Supplementary Data 2**

Summary of small RNA-seq and long RNA-seq data quality metrics for individual human oocytes.

**Supplementary Data 3**

Gene expression matrix (FPKM) for single oocytes across different developmental stages.

**Supplementary Data 4**

Expression matrix (RPM) of TE subfamilies (Sheet 1), TE-aligned short-piRNAs (Sheet 2), and TE-aligned long-piRNAs (Sheet 3) in single oocytes at different developmental stages.

**Supplementary Data 5**

Differential expression analysis of TE subfamilies between adjacent developmental stages (PF vs. PrF, GV vs. PF, MII vs. GV), as well as a direct comparison between MII and PrF stages.

**Supplementary Data 6**

Expression matrix (RPM) of miRNAs in single oocytes.

**Supplementary Data 7**

Differential expression analysis of TE-aligned short-piRNAs across adjacent stages (PF vs. PrF, GV vs. PF, MII vs. GV), and a direct comparison between MII and PrF stages.

**Supplementary Data 8**

Differential expression analysis of TE-aligned long-piRNAs across adjacent stages (PF vs. PrF, GV vs. PF, MII vs. GV), and a direct comparison between MII and PrF stages.

**Supplementary Data 9**

The top 1,000 most highly expressed piRNA clusters identified in this study and their corresponding uniquely mapped piRNA expression levels.

**Supplementary Data 10**

Genomic coverage of TE superfamilies in the top 20 most productive piRNA-generating clusters.

